# Rational design of chemically controlled antibodies and protein therapeutics

**DOI:** 10.1101/2022.12.22.521584

**Authors:** Anthony Marchand, Lucia Bonati, Sailan Shui, Leo Scheller, Pablo Gainza, Stéphane Rosset, Sandrine Georgeon, Li Tang, Bruno E. Correia

## Abstract

Protein-based therapeutics such as monoclonal antibodies and cytokines are important therapies in various pathophysiological conditions such as oncology, auto-immune disorders, and viral infections. However, the wide application of such protein therapeutics is often hindered by dose-limiting toxicities and adverse effects, namely cytokine storm syndrome, organ failure and others. Therefore, spatiotemporal control of the activities of these proteins is crucial to further expand their application. Here, we report the design and application of small molecule-controlled switchable protein therapeutics by taking advantage of a previously engineered OFF-switch system. We used Rosetta modeling suite to computationally optimize the affinity between B-cell lymphoma 2 (Bcl-2) protein and a previously developed computationally designed protein partner (LD3) to obtain a fast and efficient heterodimer disruption upon addition of a competing drug (Venetoclax). The incorporation of the engineered OFF-switch system into *α*CTLA4, anti-HER2 antibodies or an Fc-fused IL-15 cytokine demonstrated an efficient disruption in vitro, as well as fast clearance in vivo upon addition of the competing drug Venetoclax. These results provide a proof-of-concept for the rational design of controllable biologics by introducing a drug-induced OFF-switch into existing protein-based therapeutics.

## MAIN TEXT

Protein-based therapeutics, such as monoclonal antibodies (mAbs) and cytokines, have shown to mediate potent antitumor effects and are the fastest growing group of therapeutics (1-2). Nevertheless, their therapeutic use is limited by systemic toxicities arising from excessive immune and inflammatory responses, and by off-target effects (3-4). Innovative engineering strategies have been applied to increase safety through localized activity of the therapeutic (5-7) or drug-induced ON-switch system (8). However, none of these approaches allows the direct OFF-switch control of the therapeutics’ activity with an external trigger that can be applied as desired. A system that allows the spatiotemporal control of biological activities upon administration of clinically-approved small molecules represents a promising strategy to increase protein therapeutics’ safety profile. Several prior studies focused on modulating protein-protein interactions (PPIs) using small molecules to trigger either disruption or dimerization (9-13). We previously reported a novel chemically-disruptable heterodimer composed (CDH) of a BH3-motif grafted and computationally improved protein (LD3) binding to B-cell lymphoma-extra large (Bcl-XL) or B-cell lymphoma 2 protein (Bcl-2) with high affinity (14-15). The heterodimers can be disrupted by A-1155463 and Venetoclax, respectively. However, this approach has never been used to control the activity of a soluble protein therapeutic. Here, we computationally optimized the interface of the CDH for enhanced drug sensitivity and faster disruption. We used the optimized CDH to disrupt the Fc region from a therapeutic protein to control its half-life. Our results demonstrate the potential of designed OFF-switches for generating biologics with enhanced safety and broader applications.

To generate switchable antibodies (SwAbs), we placed the LD3:Bcl-2 complex, that can be disrupted by Venetoclax, between the epitope-binding region and the fragment crystallizable (Fc) region of the antibody (Fig. 1A). Fc regions are crucial for antibodies as they provide important features such as: i) longer half-life *in vivo* (16), ii) increased avidity effect due to the dimerization (17) and iii) an ability to trigger effector functions (18). We hypothesized that the addition of Venetoclax would compete for the LD3-binding site on Bcl-2 and trigger disruption between the two components. As a result, the epitope-binding domain would lose the advantages provided by the Fc-region, leading to an indirect OFF-switch of the biological activity.

**Figure 1.**
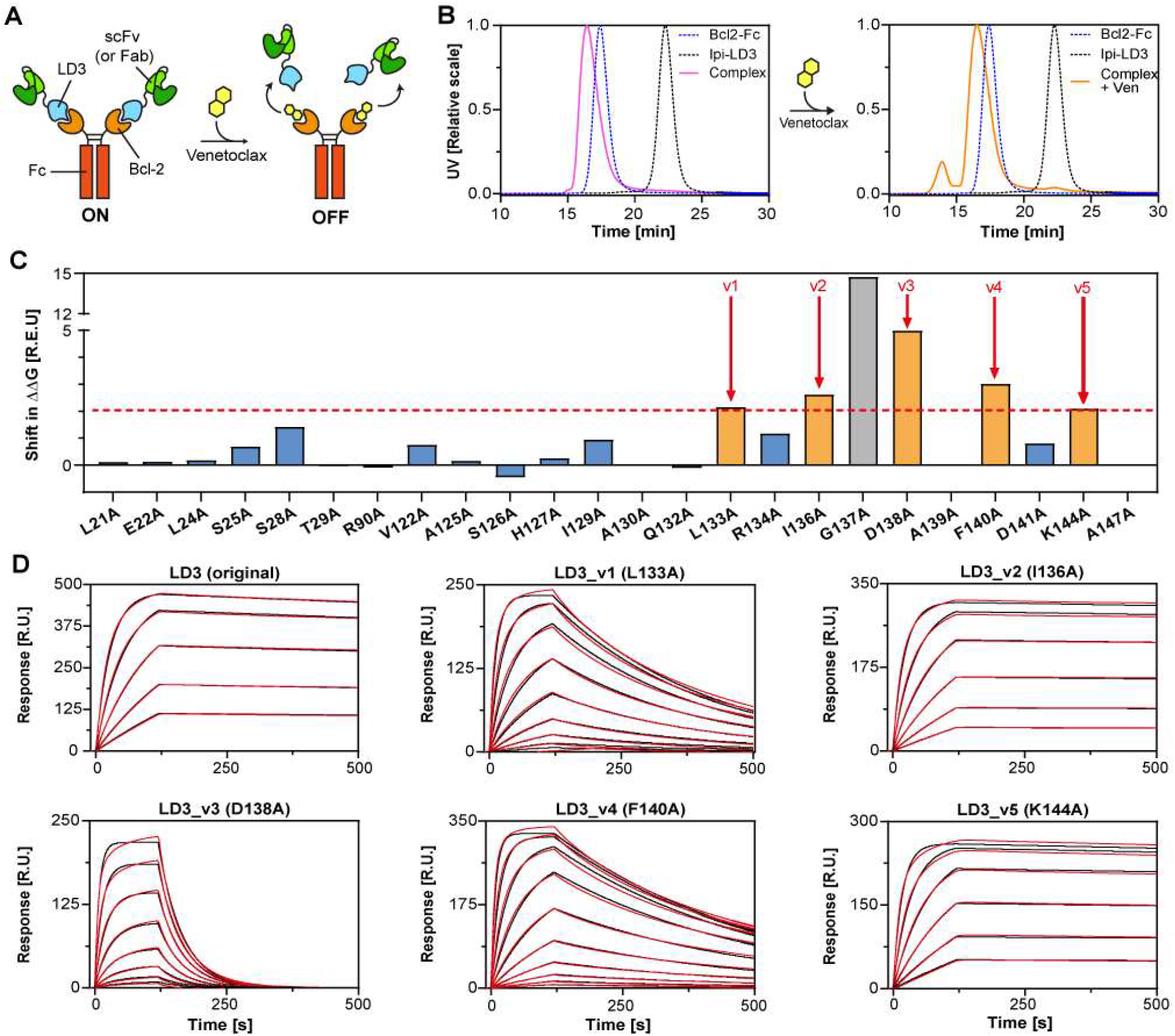
Computational design and improvement of a switchable antibody system. **A**. Schematic representation of the switchable antibody system. The single-chain variable fragment (scFv) or Fab fragment is fused to a computational design (LD3) with high affinity to the Fc-fused Bcl-2. The addition of Venetoclax binds to the LD3-binding site on Bcl-2 and triggers disruption of the switchable antibody. **B**. SEC-MALS of an *α*CTLA4 Fab fused to LD3 (Ipi-LD3, gray dashed line), a Fc-fused Bcl-2 (blue dashed line), the switchable antibody complex (pink, left part) and the switchable antibody complex incubated with Venetoclax (orange, right part). **C**. Computational alanine scan obtained with Rosetta. Mutations to alanine giving an increase of the computed binding energy (ΔΔG) of at least two Rosetta Energy Unit (R.E.U.) were considered as variant candidates (Orange bars). G137A mutation was not considered (Gray bar). **D**. Surface plasmon resonance (SPR) with Bcl-2 binding to different immobilized LD3-variants (v1 to v5). Measurements are indicated in red and fit curves in black.

We first generated a switchable version of a published *α*CTLA4 fragment antigen-binding region (Fab, Ipilimumab) (19), and tested the disruption efficiency by detecting the complex and monomeric components by size-exclusion chromatography combined with multi-angle light scattering (SEC-MALS). However, Venetoclax did not trigger detectable SwAb disruption as monomeric components were not observed (Fig. 1B, Suppl. Table 1). We therefore hypothesized that the low-nanomolar affinity of the LD3:Bcl-2 complex (Table 1) does not allow an efficient competition by the drug, most probably due to the slow dissociation rate (k_off_) that restricts the opportunity of the drug to displace the LD3 binder. With these considerations, we aimed to further engineer LD3 for reduced affinity for Bcl-2. We used the protein modeling framework Rosetta, to conduct a computational alanine-scan on all LD3 interface residues to highlight alanine mutants with increased computed binding energy (ΔΔG) (Fig.1C). All mutations to alanine increasing the ΔΔG by 2 Rosetta Energy Unit (R.E.U) were considered as potential LD3 variant candidates (v1 to v5), except for G137A which introduces a steric clash likely to be considerably deleterious for binding. The remaining five LD3 variants were expressed, purified and tested by surface plasmon resonance (SPR) for binding Bcl-2 compared to the original LD3 protein (Fig. 1C-D, Table 1 & Supp. Fig S1).

**Table 1.**
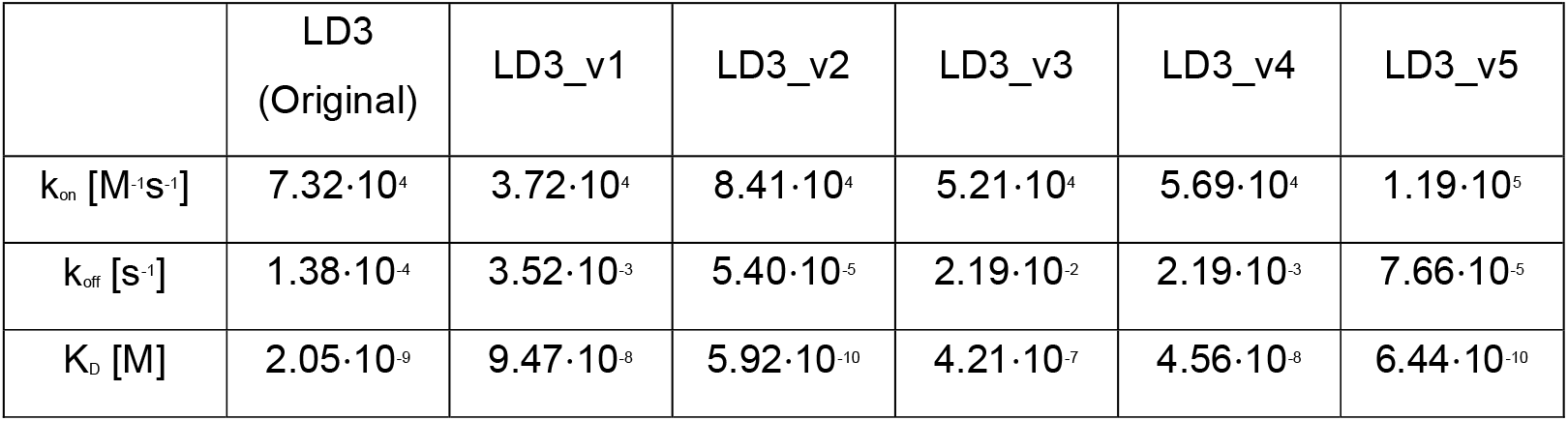
Summary table of the affinities surface plasmon resonance data of the different LD3 variants.^1^.

We sought to find variants with slightly decreased dissociation rate (k_off_) compared to the original LD3, but with unperturbed association rate (k_on_). Variants 2 (I136A) and 5 (K144A) showed only minor differences to the original LD3, and were not further considered. Variant 3 (D138A) had the highest destabilization effect, which is consistent with the high ΔΔG difference predicted by the alanine scan. Both variants 1 (L133A) and 4 (F140A) showed similar mild decreases in dissociation rates, however variant 4 had a less affected association rate and was therefore chosen as a lead candidate for the switchable antibody system. We used LD3 variant 4 (LD3_v4) to generate an improved version of the switchable Ipilimumab-based *α*CTLA4 antibody by fusing the Ipilimumab Fab to LD3_v4. After complex formation with Bcl2-Fc, we assessed the switchability using SEC-MALS as described above. While only 3% (muncomplex/mtotal) of the switchable antibodies were disrupted on SEC-MALS upon Venetoclax treatment with the original LD3 protein, more than 90% of the complex efficiently disrupted with LD3_v4 (Fig. 2A, Suppl. Table 1). We evaluated disruption kinetics by biolayer interferometry (BLI) and detected 30% disruption at the highest tested concentration of Venetoclax (10 *μ*M) after 200 seconds (Fig. 2B). During that time, the switchable antibody complex remained stable in solution without addition of Venetoclax. To confirm these results in a cell-based assay, we substituted the antigen-targeting domain of the SwAb with an *α*HER2 scFv that allowed the labeling of HER2-expressing cells.

**Figure 2.**
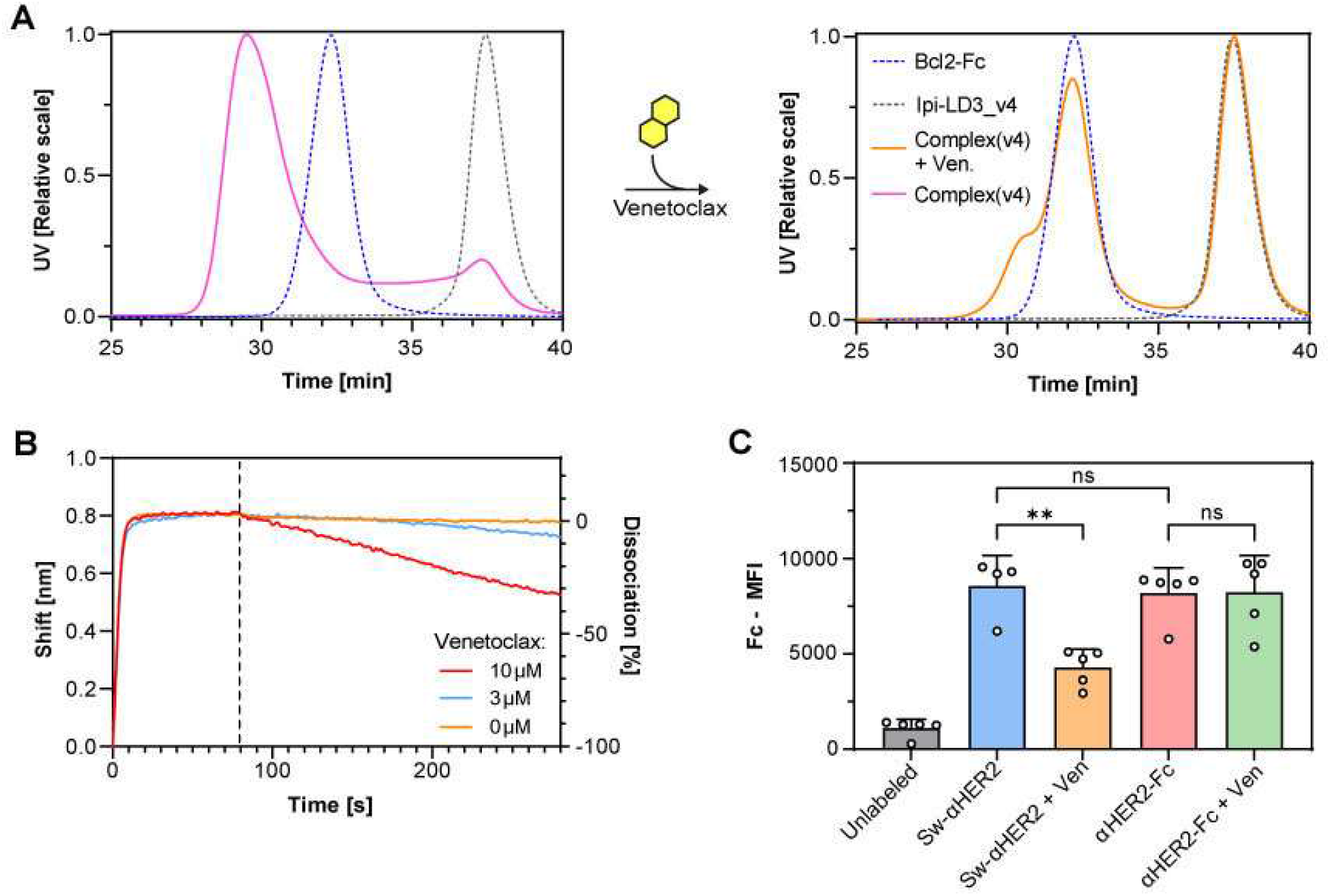
Disruption efficiency of a switchable antibody with LD3_v4. **A**. SEC-MALS of a Bcl2-Fc alone (blue dashed line), an *α*CTLA4 Fab (Ipilimumab) fused to LD3 variant 4 (gray dashed line) and the switchable antibody complex in absence (pink) and presence (orange) of Venetoclax. **B**. Biolayer interferometry (BLI) measurements of the switchable anti-CTLA4 antibody with increasing concentration of Venetoclax. **C**. Quantification of the mean fluorescence intensity (MFI) measured on the surface of MC38 cells unlabeled or labeled with with a switchable or conventional *α*Her2 antibody (Sw-*α*HER2 and *α*HER2-Fc respectively) treated with and without Venetoclax. Tukey’s multiple comparisons test, p<0.01 (**), non-significant (ns).

We stained MC38-HER2 cells, a murine colon adenocarcinoma cell line stably expressing HER2, with the *α*HER2 SwAb and treated the cells with or without Venetoclax (Fig. 2C and Supp. Fig. S2). One hour after adding Venetoclax, the Fc fragment detected on MC38 cell surface decreased by 2-fold. Among other possibilities, the reduced Venetoclax-induced antibody disruption might be explained by the avidity provided by the two Fabs binding simultaneously, which may reduce drug sensitivity. The switchable *α*HER2 antibody showed similar binding to MC38-HER2 cells compared to a conventional *α*HER2 antibody, which did not respond to Venetoclax. Altogether, these results confirm the improved switchability of the engineered antibody.

We next tested the function of the engineered switchable proteins *in vitro* and *in vivo* by measuring cell proliferation and the half-life in mice blood. To do so, we extended the strategy to the generation of switchable cytokines. We chose mouse IL-15 superagonist (IL-15SA) and generated switchable IL-15SA (SwIL-15SA) by fusing IL-15 and the IL-15 receptor *α* domain (IL-15R*α*) to LD3 assembled with Bcl2-Fc (Fig. 3A). To assess the functionality of SwIL-15SA, we stimulated primary murine T cells *ex vivo* with either IL-15SA or SwIL-15SA and measured cell proliferation.

**Figure 3.**
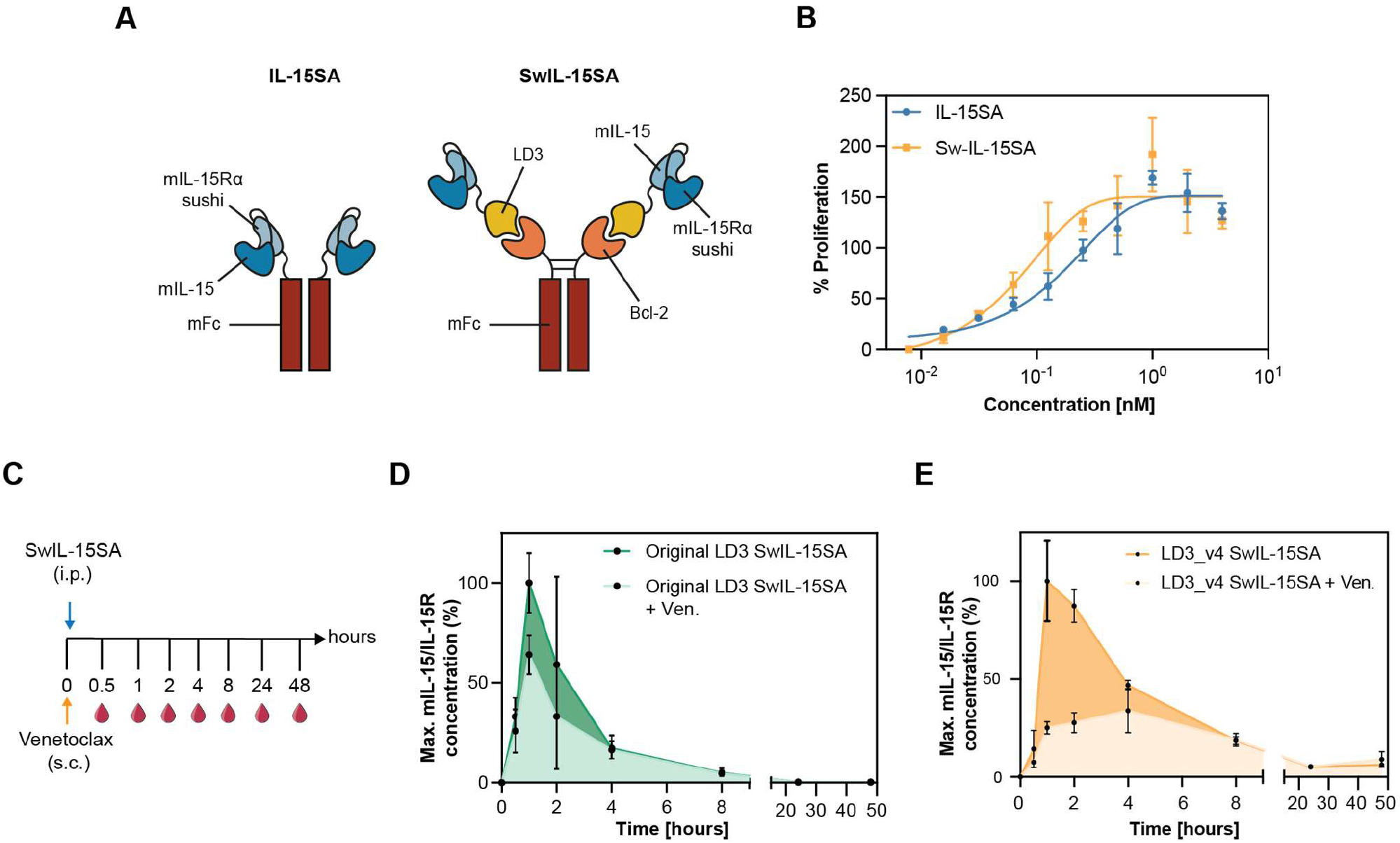
Functional assessment and *in vivo studies* using an Fc-fused switchable cytokine. **A**. Schematic representation of the switchable interleukin system. In IL-15SA, the sushi domain of mouse IL-15R*α* is fused to mIL-15 binding a mousse Fc (left). In SwIL-15SA, the sushi domain of mouse IL-15R*α* is fused to the optimized LD3 binding to mouse Fc-fused Bcl-2 (right). **B**. Activated mouse T cell proliferation in response to IL-15SA or SwIL-15SA. **C**. C57BL/6 mice were first injected subcutaneously (s.c.) with Venetoclax (25.0 mg/kg) and subsequently injected intraperitoneally (i.p.) with 100 pmol SwIL-15SA. Mice were bled overtime after 0.5, 1, 2, 4, 8, 24, and 48 hours after treatment. **D**. Pharmacokinetic properties of SwIL-15SA composed of the IL-15/IL-15R complex fused to the original LD3 with (light green) or without (dark green) the administration of Venetoclax. **E**. Pharmacokinetic properties of SwIL-15SA composed of the IL-15/IL-15R complex fused to LD3_v4 with (light orange) or without (dark orange) the administration of Venetoclax.

Proliferation of murine primary T cells induced by SwIL-15SA was comparable to conventional IL-15SA, indicating that fusing LD3 to the sushi domain of IL-15R*α* did not hinder its functionality (Fig. 3B). In a second step, we assessed the switchability of SwIL-15SA *in vivo*. C57BL/6 mice were first injected subcutaneously (s.c.) with or without Venetoclax and then intraperitoneally (i.p.) with SwIL15-SA containing the original LD3. Mice were bled overtime after treatment and IL-15/IL15R complex concentration was measured by enzyme-linked immunosorbent assay (ELISA) (Fig. 3C). IL-15/IL-15R complex concentration in blood of mice treated with Venetoclax peaked at 60% of the maximum IL-15/IL15R concentration of the control group, suggesting that Venetoclax administration induced a disruption of the Bcl-2:LD3 complex, which led to fast clearance of the IL-15-derived domains (Fig. 3D).

To investigate whether the affinity of the Bcl-2:LD3 complex could provide an additional parameter to tune the switchability efficiency of the system, we further tested a variant of SwIL-15SA being composed of the IL-15 sushi domain fused to LD3_v4. Here, blood concentrations in control mice peaked at 1 hour after injection and then decreased overtime (Fig. 3E). Unlike the control group, in mice treated with Venetoclax the IL-15/IL15R complex concentration reached only about 25% of the maximum IL-15/IL15R concentration of the control group. This observation suggests that the disruption efficiency and the half-life of the system can be tuned with the affinity of the Bcl-2:LD3 complex. Overall, these results show that Venetoclax disrupts the interaction between Bcl-2 and LD3, leading to the fast clearance of monomeric IL-15/IL-15R-LD3 *in vivo*.

Altogether, we show a modular and generalizable OFF-switch approach for the design of safe antibody and cytokine therapeutics by introducing a chemically-disruptable heterodimer between the therapeutic domain and the Fc moiety. Loss of the Fc-fragment leads to a decrease of the avidity effect and a drastic reduction of the protein half-life. We took advantage of a previously designed CDH that can be competed by a clinically-approved drug, Venetoclax, which makes it a good candidate for translational applications. Of note, one strength of our system is its modularity with the ease to adapt it to several therapeutic proteins by exchanging the therapeutic domain fused to LD3. But the large size of the protein complex (of about ∼250 kDa for a switchable antibody, compared to ∼150 kDa for a normal antibody) may limit tissue penetration (20). However, for highly toxic therapies, such as immunostimulatory therapies, these limitations would be outweighed by the improved safety profile. Our presented workflow to reduce heterodimer affinity to increase drug sensitivity can likely be readily extended to other examples of CDHs. These types of switchable biologics could serve as a basis for safer biologics for therapeutic use.

## AUTHOR INFORMATION

## Author Contributions

Anthony Marchand‡, Lucia Bonati‡, Li Tang and Bruno E. Correia leaded the project. Anthony Marchand‡, Lucia Bonati‡ performed the experimental work. Sailan Shui, Leo Scheller and Pablo Gainza contributed to design the experimental set-up. Anthony Marchand‡ and Pablo Gainza contributed to the computational optimization. Stéphane Rosset performed the bio-layer interferometry. Sandrine Georgeon expressed and purify the proteins. Anthony Marchand‡, Lucia Bonati‡, Li Tang and Bruno E. Correia wrote the manuscript with input from all authors. ‡ These authors contributed equally.

## Notes

The authors declare that a patent has been filed describing the findings reported in the paper.

## ACKNOWLEDGMENT

We thank the EPFL animal facility (CPG), for their support for conducting animal experiments, and the flow cytometry core facility (FCCF) for their assistance. We also thank the high-performance computing facility at EFPL – SCITAS for the computational resources.

## Footnotes

1 Data was collected using SPR showing the association rate (k_on_), dissociation rate (k_off_) and dissociation constant (KD) of the original LD3 and the different variants obtained by computational alanine scanning.

