## Supplementary material for "Rational design of chemically controlled antibodies and protein therapeutics": Material and Methods

*Computational design*

Previously solved crystal structure of Bcl-2 in complex with LD3 was used for computational modeling (PDB ID: 6IWB). Using the Rosetta modeling suite, the pose was relaxed with the "FastRelax" mover, before the computational alanine scan was performed using an "Alascan" filter. Residues where a mutation to alanine lead to an increase of the computed binding energy of >2 Rosetta energy units (R.E.U.) were considered as potential candidates to lower the affinity of LD3 for Bcl2. Mutations exceeding 5 R.E.U. were not considered due to the introduction of clashes that may abrogate binding.

*Protein expression and purification*

The engineered IL-15SA construct (gWIZ-mIL-15SA) was a gift from D. J. Irvine (MIT). IL-15SA contains a mouse IL-15 fused at the C terminus of Sushi domain of a mouse IL-15R$\alpha$, which is next fused at the C terminus with a mouse IgG2c Fc. A previously optimized version of Bcl-2 (1) was fused to either human IgG or mouse IgG2 Fc-fragment (see supplementary table 2). Switchable antibodies were composed of either a previously published $\alpha$CTLA4 antibody (2) Ipilimumab as a Fab or an $\alpha$HER2 4D5 clone (3) as an scFv fused to LD3 protein N-terminal with a $(GGGS)_3$-linker. As a switchable cytokine, we used a fusion protein composed of mouse IL-15 C-terminally fused to the IL-15 receptor $\alpha$ domain (IL-15R$\alpha$), itself fused to the LD3 variant C-terminal with a $(GGGS)_3$-linker. DNA sequences were ordered from Twist Bioscience and Gibson cloning used to clone into bacterial (pET11) or mammalian (pHLSec) expression vectors. Mammalian expressions were performed using the Expi293$^{TM}$ expression system from Thermo Fisher Scientific. Supernatant was collected 6 days post transfection, filtered, and purified. *E. coli* expressions were performed using BL21 (DE3) cells and IPTG induction (1 mM at OD 0.6-0.8) and growth overnight at 16-18° C. Pellets were lysed in lysis buffer (50 mM Tris, pH 7.5, 500 mM NaCl, 5% Glycerol, 1 mg/ml lysozyme, 1 mM PMSF, and 1 µg/ml DNase) with sonication, the lysate clarified, and purified. Proteins were then purified using an ÄKTA pure system (GE healthcare) with Ni-NTA affinity columns followed by size exclusion chromatography with PBS.

*Surface plasmon resonance*

SPR measurements were performed on a Biacore 8K (GE Healthcare) with HBS-EP+ as running buffer (10 mM HEPES pH 7.4, 150 mM NaCl, 3 mM EDTA, 0.005% v/v Surfactant P20, GE Healthcare). Original LD3 and mutants were immobilized on a CM5 chip (GE Healthcare # 29104988) via amine coupling. 500-1000 response units (RU) were immobilized and Bcl-2 was injected as an analyte in serial dilutions. The flow rate was 30 µl/min for a contact time of 120s followed by 400s dissociation time. After each injection, the surface was regenerated using 50 mM NaOH. SPR Data were fit with 1:1 Langmuir binding model within the Biacore 8K analysis software (GE Healthcare #29310604).

*Bio-layer interferometry (BLI)*

Measurements were performed on a Gator BLI system. The running buffer was PBS. Fc-tagged Bcl-2 were diluted to 5 $\mu$g/mL and immobilized on the anti-human IgG tips for 80 seconds (1-2 nm immobilized). The loaded tips were then dipped into 500 nM LD3-fused Ipilimumab Fab (or PBS for the reference) for 80 seconds and then in different concentrations of Venetoclax (10, 3 and 0 $\mu$M) diluted in PBS for 210 seconds. Each measurement was subtracted with the reference (channel with Fc-fused Bcl2 immobilized, no associated LD3 and a corresponding concentration of Venetoclax diluted in PBS).

*Size exclusion chromatography multi-angle light scattering (SEC-MALS)*

Size exclusion chromatography with an online multi-angle light scattering device (miniDAWN TREOS, Wyatt) was used to determine the oligomeric state and molecular weight for the switchable antibodies in solution. Purified LD3-Fab and Bcl2-Fc proteins were mixed with a 2:1 molar ratio and incubated at room temperature for 5 min to form a complex. Assembled complexes received 100 $\mu$M Venetoclax or PBS and incubated 1h at 37°C. Final concentration was approximately 1 mg/ml in PBS (pH 7.4), and 100 µl of the sample was injected into a Superdex 75 300/10 GL column (GE Healthcare) with a flow rate of 0.5 ml/min, and UV280 and light scattering signals were recorded. Molecular weight was determined using the ASTRA software (version 6.1, Wyatt).

*In vitro cell binding assay*

100’000 HER2-transduced MC38 mouse colon cancer cells were collected in a tube. Purified HER2-specific LD3-Fab and Bcl-2-Fc proteins were mixed at a 2:1 ratio and incubated at room temperature for 5 minutes to form a complex. MC38-HER2+ cells were then stained with $\alpha$HER2 SwAb at concentrations of 100 nM and incubated at 4 °C for 30 min. An Fc-fused $\alpha$HER2 ($\alpha$HER2_Fc) was used as a positive control. Cells were washed twice with FACS buffer (PBS containing bovine serum albumin, 0.2% (w/v)) and 10 µM Venetoclax was added to the cells and incubated at 37 °C for 1 hour. Following, cells were washed and stained with anti-human Fc antibody at 4 °C fr 30 min. Cells were then washed, stained with 4′,6-diamidino-2-phenylindole (DAPI; Sigma-Aldrich) and analyzed by FACS.

*T-cell proliferation assay*

Activated Pmel T cells were collected by centrifugation, re-suspended in mouse T-cell media and seeded at a density of 10’000 T cells/well in a 96-well flat bottom tissue culture plate. T cell growth was stimulated by addition of serial dilutions of IL-15SA or Sw-IL15SA to a total volume of 100 µL and cultured for 48 hours at 37 °C. On day 2, cells were collected, washed once with FACS buffer and stained with DAPI. Cell counts for each condition were quantified by FACS using the Attune NxT flow cytometer (Invitrogen/Thermo Fisher Scientific).

*Animal studies*

6-8 week-old female C57BL/6 mice were purchased from Charles River Laboratories and maintained in the animal core facility [Center of Phenogenomics (CPG)] of École Polytechnique Fédérale de Lausanne (EPFL). In evaluating the switchability potential of SwIL-15SA, C57BL/6 mice were injected subcutaneously with 100 µl Venetoclax dissolved at 25 mg/kg in a solution of saline and 2% dimethyl sulfoxide (DMSO). Following, the animals were injected intraperitoneally with 100 pmol of Sw-IL15SA in 100 µl and bled overtime at 0.5, 1, 2, 4, 8, 24, and 48 hours after treatment. IL-15/IL-15R complex concentration in blood was quantified using a commercial enzyme-linked immunosorbent assay (ELISA) kit following manufacturer's instructions (Thermo Fisher Scientific, 88-7215-88).
