## Supplementary Table 1 for "Rational design of chemically controlled antibodies and protein therapeutics"

| | Mass fraction [%] | | |
|---|---|---|---|
| | Complex | Bcl2-Fc | Ipi-LD3 |
| Original LD3 | 97.0% | 3% | N/D |
| LD3_v4 | 9.6% | 36% | 54.2% |

**Supplementary Table 1: Mass fraction of the different SwAb components measured by the SEC-MALS upon Venetoclax treatment**. Switchable antibody complexes were assembled *in vitro* with either the original LD3 or the variant 4 (LD3_v4), treated with venetoclax, and analyzed by size-exclusion chromatography coupled to multi-angle light scattering (SEC-MALS). The mass fraction of the different peaks constituting the disrupted complex were measured.
