## Supplementary Table 2 for "Rational design of chemically controlled antibodies and protein therapeutics"

| | |
|---|---|
| Bcl2-hmFc | AHAGRTGYDNREIVMKYIHYKLSQRGYEWDAGDDAEENRTEAPEGTESEVVHRALRD<br>AGDDFERRYRRDFAEMSSQLHLTPDTARQRFETVVEELFRDGVNWGRIVAFFEFGGV<br>MCVESVNREMSPLVDNIAEWMTEYLNRHLHTWIQDNGGWDAFVELYGPSMRGGGG<br>SGTDKTHTCPPCPAPELLGGPSVFLFPPKPKDTLMISRTPEVTCVVVDVSHEDPEVKF<br>NWYVDGVEVHNAKTKPREEQYNSTYRVVSVLTVLHQDWLNGKEYKCKVSNKALPAPI<br>EKTISKAKGQPREPQVYTLPPSREEMTKNQVSLTCLVKGFYPSDIAVEWESNGQPEN<br>NYKTTPPVLDSDGSFFLYSKLTVDKSRWQQGNVFSCSVMHEALHNHYTQKSLSLSPG<br>KHHHHHH |
| Bcl2-mFc | AHAGRTGYDNREIVMKYIHYKLSQRGYEWDAGDDAEENRTEAPEGTESEVVHRALRD<br>AGDDFERRYRRDFAEMSSQLHLTPDTARQRFETVVEELFRDGVNWGRIVAFFEFGGV<br>MCVESVNREMSPLVDNIAEWMTEYLNRHLHTWIQDNGGWDAFVELYGPSMRGGGG<br>SEPRVPITQNPCPPLKECPPCAAPDLLGGPSVFIFPPKIKDVLMISLSPMVTCVVVAVSE<br>DDPDVQISWFVNNVEVHTAQTQTHREDYNSTLRVVSALPIQHQDWMSGKEFKCKVN<br>NRALPSPIEKTISKPRGPVRAPQVYVLPPPAEEMTKKEFSLTCMITGFLPAEIAVDWTS<br>NGRTEQNYKNTATVLDSDGSYFMYSKLRVQKSTWERGSLFACSVVHEGLHNHLTTKT<br>ISRSLGKGTKHHHHHH |
| LD3 | GQRWELALGRFLEYLSWVSTLSEQVQEELLSSQVTQELRALMDETMKELKAYKSELE<br>EQLTPVAEETRARLSKELQAAQARLGADMEDVRGRLVQYRGEVQAMLGQSTEELRV<br>RLASHLIALQLRLIGDAFDLQKRLAVYQAGA |
| Ipilimumab_H | QVQLVESGGGVVQPGRSLRLSCAASGFTFSSYTMHWVRQAPGKGLEWVTFISYDGN<br>NKYYADSVKGRFTISRDNSKNTLYLQMNSLRAEDTAIYYCARTGWLGPFDYWGQGTL<br>VTVSSASTKGPSVFPLAPSSKSTSGGTAALGCLVKDYFPEPVTVSWNSGALTSGVHTF<br>PAVLQSSGLYSLSSVVTVPSSSLGTQTYICNVNHKPSNTKVDKRVEPKSC |
| Ipilimumab_L | EIVLTQSPGTLSLSPGERATLSCRASQSVGSSYLAWYQQKPGQAPRLLIYGAFSRATG<br>IPDRFSGSGSGTDFTLTISRLEPEDFAVYYCQQYGSSPWTFGQGTKVEIKRTVAAPSV<br>FIFPPSDEQLKSGTASVVCLLNNFYPREAKVQWKVDNALQSGNSQESVTEQDSKDST<br>YSLSSTLTLSKADYEKHKVYACEVTHQGLSSPVTKSFNRGEC |
| mIL15 | GTTCPPPVSIEHADIRVKNYSVNSRERYVCNSGFKRKAGTSTLIECVINKNTNVAHWTT<br>PSLKCIRDPSLAGGSGGSGGSGGSGGSGGSGGNWIDVRYDLEKIESLIQSIHIDTTLYT<br>DSDFHPSCKVTAMNCFLLELQVILHEYSNMTLNETVRNVLYLANSTLSSNKNVAESGC<br>KECEELEEKTFTEFLQSFIRIVQMFINTSHHHHHH |
| $\alpha$HER2_scFv | EVQLVESGGGLVQPGGSLRLSCAASGFNIKDTYIHWVRQAPGKGLEWVARIYPTNGY<br>TRYADSVKGRFTISADTSKNTAYLQMNSLRAEDTAVYYCSRWGGDGFYAMDYWGQG<br>TLVTVSSGGGGSGGGGSGGGGSDIQMTQSPSSLSASVGDRVTITCRASQDVNTAVA<br>WYQQKPGKAPKLLIYSASFLYSGVPSRFSGSRSGTDFTLTISSLQPEDFATYYCQQHY<br>TTPPTFGQGTKVEIK |
| $\alpha$HER2_Fc | DYKDIVMTQSPSSLSASVGDRVTITCRASQDVNTAVAWYQQKPGKAPKLLIYSASFLY<br>SGVPSRFSGSRSGTDFTLTISSLQPEDFATYYCQQHYTTPPTFGQGTKVELKRATPSH<br>NSHQVPSAGGPTANSGEVKLVESGGGLVQPGGSLRLSCATSGFNIKDTYIHWVRQAP<br>GKGLEWVARIYPTNGYTRYADSVKGRFTISADTSKNTAYLQMNSLRAEDTAVYYCSR<br>WGGDGFYAMDYWGQGTTVTVSSTGVHSEPRVPITQNPCPPLKECPPCAAPDLLGGP<br>SVFIFPPKIKDVLMISLSPMVTCVVVAVSEDDPDVQISWFVNNVEVHTAQTQTHREDYN<br>STLRVVSALPIQHQDWMSGKEFKCKVNNRALPSPIEKTISKPRGPVRAPQVYVLPPPA<br>EEMTKKEFSLTCMITGFLPAEIAVDWTSNGRTEQNYKNTATVLDSDGSYFMYSKLRVQ<br>KSTWERGSLFACSVVHEGLHNHLTTKTISRSLGKASGSRSLLANKRSEL |

**Supplementary table 2: Amino acids sequences of the different proteins used**. A stabilized version of Bcl2 fused to either human IgG1 (Bcl2-hmFc) or mouse IgG2 (Bcl2-mFc), a previously defined Bcl2-binding protein (LD3), an anti-CTLA4 Fab (Ipilimumab_H and Ipilimumab_L), a mouse interleukin 15 (mIL15), an anti-HER2 single chain fragment clone 4D5 ($\alpha$HER2_scFv) and the same fused to an IgG ($\alpha$HER2_Fc).
