## Supplementary figures and images for "Rational design of chemically controlled antibodies and protein therapeutics"

### Supplementary Figure S1

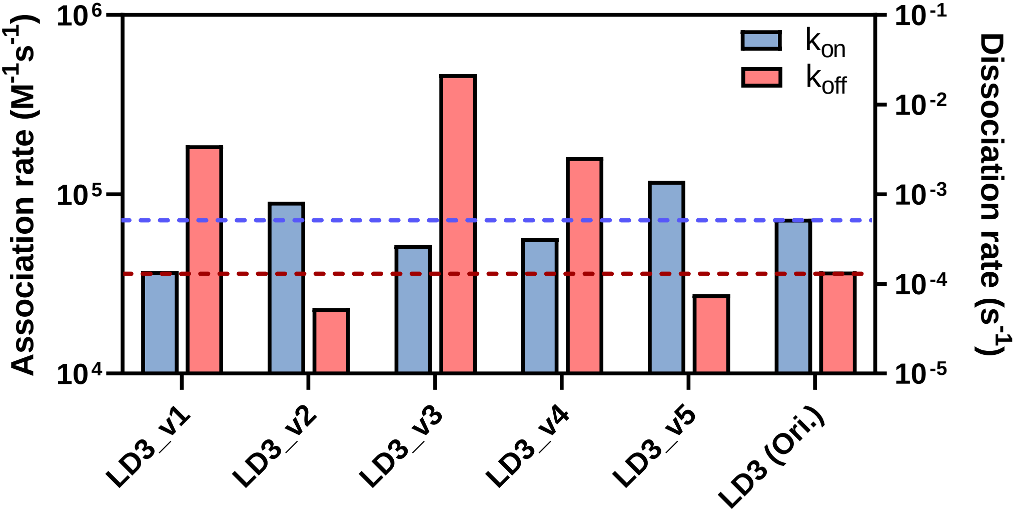

### Supplementary Figure S2

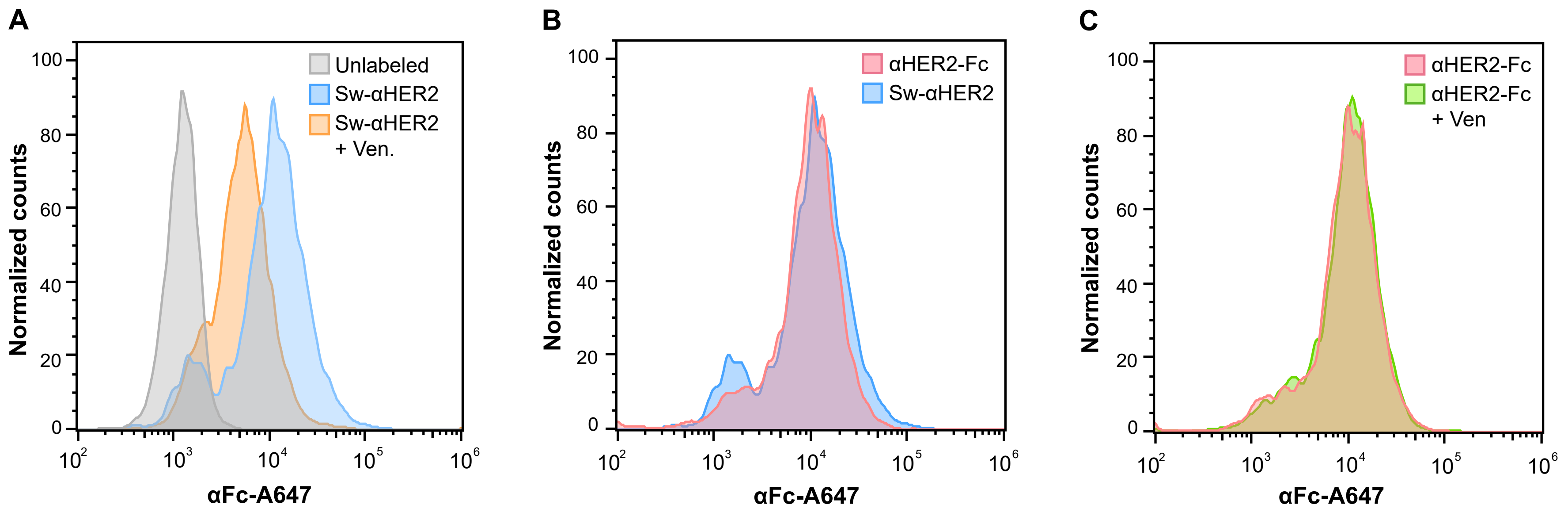
